# Logical design of synthetic cis-regulatory DNA for genetic tracing of cell identities and state changes

**DOI:** 10.1101/2022.11.04.515171

**Authors:** Carlos Company, Matthias Jürgen Schmitt, Yuliia Dramaretska, Sonia Kertalli, Ben Jiang, Michela Serresi, Iros Barozzi, Gaetano Gargiulo

## Abstract

Descriptive data are rapidly expanding in biomedical research. Instead, functional validation methods with sufficient complexity remain underdeveloped. Transcriptional reporters allow experimental characterization and manipulation of developmental and disease cell states, but their design lacks flexibility. Here, we report logical design of synthetic *cis*-regulatory DNA(LSD), a computational framework leveraging phenotypic biomarkers and *trans*-regulatory networks as input to design reporters marking the activity of selected cellular states and pathways. LSD uses bulk or single-cell biomarkers and a reference genome or custom *cis*-regulatory DNA datasets with user-defined boundary regions. By benchmarking validated reporters, we integrated LSD with a computational classifier to rank phenotypic specificity of putative *cis*-regulatory DNA. Experimentally, LSD-designed reporters targeting a wide range of cell states are functional without minimal promoters. *In silico*, an LSD-unsupervised mesenchymal glioblastoma reporter outperformed previously validated ones. In genome-scale CRISPRa screens, it discovered known and novel *bona fide* cell-state-drivers. Thus, LSD captures core principles of *cis*-regulation and is broadly applicable to studying complex cell states and mechanisms of transcriptional regulation.

## Introduction

The precise identification of specific cell types and transient states is essential to understanding biological processes in which a diverse set of cell types/states contributes to tissue homeostasis. Accurately defining cell entities, states, as well as boundaries and trajectories governing physiological and pathological transitions is particularly important for understanding cell responses to complex alterations of homeostasis, such as cancer^1-3^ and viral infections ^4^. Moreover, monitoring the spatiotemporal activation of a given pathway is instrumental to dissecting the underlying biology as well as monitoring the response to biological and chemical perturbations. While single-cell genomics and proteomics are providing increasingly powerful multiomics maps of cellular processes in steady-state conditions and in longitudinal analyses, equally comprehensive experimental tools to trace live cells are lagging behind. Lineage tracing in developmental settings exploits genetic tagging of a single gene map the fate of phenotypes associated with the expression of the selected gene^5^. Limitations associated with engineering an endogenous locus with a reporter include the assumption that gene expression regulation of the selected biomarker is a direct proxy of the phenotype of interest. This may not be systematically warranted when complex regulatory netoworks are studied. Conversely, selecting *cis-*regulatory elements to design synthetic cassettes showing sufficient specificity is complicated by our incomplete functional annotation and the mechanistical understanding of *cis*-regulation for most genes.

Synthetic transcriptional reporters may be assembled by juxtaposing candidate *cis-*regulatory DNA sequences. In cellular and molecular genetics, designing synthetic reporters starting from naturally occurring *cis-*regulatory elements responsive to well-defined signaling pathways or to combinations of transcription factors is a well-established strategy^6,7^. Significant effort was directed towards generating and selecting synthetic reporters using massively parallel sequencing or mixed computational design strategies^8-11,12^. This revealed the outstanding potential of this approach, as well as the limitations associated with incomplete control over the design, which remains challenging^9,11^. Importantly, how biochemically defined endogenous *cis*-regulatory DNA informs the generation of synthetic enhancers remains unclear ^13^. As a rule of thumb, success in generating functional reporters is dramatically increased by combining candidate enhancers with validated *cis-*regulatory elements, such as viral or the β-globin minimal promoter. This, however, holds undefined consequences over the specificity and sensitivity of such reporters.

We recently developed a method to generate synthetic transcriptional reporters for genetic tracing (termed synthetic locus control region, or sLCR) and used these to study the significance of glioblastoma heterogeneity *in vitro* and *in vivo*^14^. This method is potentially applicable to a variety of biological settings when a more streamlined, automated implementation of computational workflow is implemented. Here, we present a computational framework to *de novo* assemble functional sLCRs capable of working on stereotyped inputs and returning an optimal candidate sLCR output. Logical design of synthetic *cis-*regulatory DNA (or LSD) generates one candidate sLCR from a user-dependent list of biomarkers and transcription factors by performing an unbiased search for optimal cis-regulatory DNA within the reference genome, or a user-defined set of candidate *cis-*regulatory elements. We complement LSD with a computational approach to rank candidates with naturally occurring or synthetic DNA based on the signal-to-noise ratio of a phenotype of interest. In turn, we benchmarked LSD’s performance using validated reporters and offered a proof-of-concept on how to exploit LSD towards the systematic characterization of cell types and states as well as to validate bulk and single-cell genomic studies.

## Results

### Logical design of synthetic *cis-*regulatory DNA

To make the design of sLCRs robust and generally applicable, we developed a fully automated framework that couples the selection of putative phenotype-specific *cis-*regulatory elements (CREs) to an iterative ranking in descending order of phenotypic representation.

The novel computational framework, termed logical design of synthetic *cis-*regulatory DNA or LSD, uses two inputs: (1) a list of *signature genes*, which are biomarkers representative of the target phenotype, and (2) a list of transcription factors (TF) with known DNA-binding motifs potentially regulating such genes (**Methods**). Both lists can be based on differential or absolute gene expression, but in principle could be defined based on any set of criteria (**Fig. 1a**). Building on our first generation sLCR design algorithm^14^, LSD first scans the regulatory landscapes of the signature genes to predict putative *cis-*regulatory elements (CREs) regulating them (**Fig. 1a** and **Methods**). By default, the boundaries of these regulatory landscapes are defined using annotated CTCF binding sites^15,16^ flanking the signature genes. Such ‘nearest CTCF neighbor’ criterion conservatively approximates the functional definition of chromatin loops ^17^ and topologically associated domains ^18^. Alternatively, users can impose experimentally-defined boundary regions, including from ChIP-seq data for other structural DNA-binding proteins and chromosome conformation capture experiments (see below). LSD scans these regulatory landscapes using a 150 bp sliding window, by 50 bp steps. This process returns a pool of putative CREs that is scored and ranked using the set of TF-binding models defined by the user. The scoring uses the following criteria: (*I*) the absolute number of TFBS; (*II*) the diversity of the TFs showing high affinity for the regions; and (*III*) the distance from the nearest endogenous transcriptional start site (TSS) (**Fig. 1a** and **Methods**). Candidate sLCRs are then generated from these CREs. The goal is to include the smallest number of CREs that faithfully represent the complexity of the regulatory inputs. To do so, LSD iteratively ranks the highest scoring CREs until all pre-defined TFs showing at least one binding site are represented (**Fig. 1a** and **Methods**). The output of LSD is proportional to the number of input TFs, the number of signature genes and the size of the genomic loci containing these genes. While our implementation relies on the scheme and assumptions outlined above, our dynamic strategy can be complemented or replaced by one based on a different set of rules or defined by hypothesis-driven criteria, such as focusing on user-defined TFBS representation.

**Figure 1.**
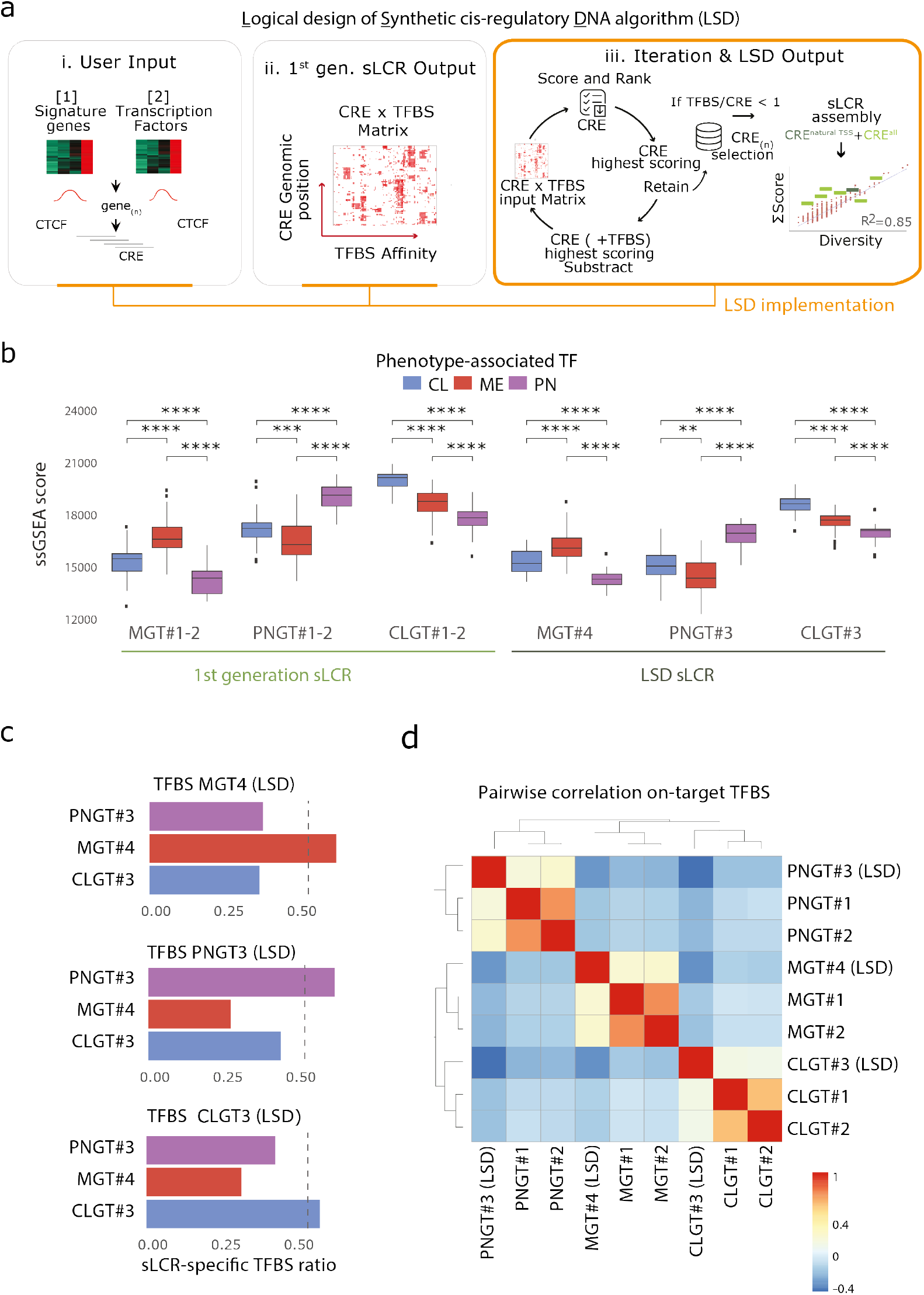
LSD streamlines the design of sLCRs from defined inputs. a) Schematic depiction of the LSD pipeline: from input signature genes and TFBS lists (i) it generates a CRE x TFBS matrix (ii; see Methods) and performs iterative selection of the top-ranked CREs (iii). Each iteration removes the highest scoring CRE and TFBS from the CRE x TFBS matrix until the CRE x TFBS contained no TFBS or CRE. The output of LSD is a ranked list of n CREs. The CRE closest to a natural TSS is prioritized. The example to the right illustrates a linear relationship between TFBS affinity and TFBS diversity for all CREs in the CRE x TBFS matrix (red circles; R2 = 0.86). In light green boxes, LSD ranked n=7 CREs (1050bp) covering >60% of the TFBS diversity. The TSS-containing CRE is in dark green. b) Boxplot showing ssGSEA scores of the TCGA Glioblastoma subtype-specific TF input lists. Each annotated GBM transcriptional subtype TF gene set (CL – Classical, blue; MES – Mesenchymal, red; PN – Proneural, purple) features comparisons by Wilcox.test (padj<0.05). c) Barplot showing the coverage of sLCR-specific TFBS lists (color) relative to the indicated TFBS input list (above). The dashed line denotes a threshold of 50%. d) Heatmap showing the Pearson correlation between the TFBS score/diversity for each sLCRs-input TF list.

To directly compare reporters generated through the first-generation approach^14^ to those designed with LSD, we designed three novel reporters for recurrent glioblastoma expression subtypes. The PNGT#3, CLGT#3 and MGT#4 sLCRs were designed in an unbiased manner by LSD and their specific signature genes were identical to those of the first-generation sLCRs (**Fig. s1a-b**), while we defined the TF lists by differential enrichment (see **Methods**). A minor variation in the TFBS list permits the design of different reporters to potentially target the same phenotype. Single-sample gene set enrichment analysis (ssGSEA) of either TF list in TCGA GBM RNA-seq data showed that they are both representatives of their target phenotype (**Fig. 1b and Supplementary Table s1**). Of note, each reporter significantly enriched the TFBS sites specific for the intended glioblastoma subtype (**Fig 1c and s1c-d**). This indicates that LSD maintains the robustness of the original approach, while operating fully automated. Importantly, hierarchical clustering of TFBS enrichment in reporters generated by either algorithm showed that the mesenchymal glioblastoma-subtype sLCR designed by LSD clustered with previously validated mesenchymal reporters, in all the tested analyses (**Fig. 1d and s1**).

### LSD allows for designing functional and specific sLCRs

To assess the performance of the LSD method, we next synthesized the LSD-designed mesenchymal glioblastoma sLCR (hereupon ‘MGT#4’) and tested it head-to-head against first-generation MGT#1-2 sLCRs constructed by user-supervised assembly and experimentally validated *in vitro* and *in vivo*^14^. Multiple sequence alignment of the three sLCRs shows only one instance of conserved positional overlap for contiguous nucleotides of the size of a TFBS (>5bp; **Fig. 2a**). This suggests that the TFBS grammar is sLCR-specific, despite all reporters targeting the same phenotype through transcription factor binding. Nevertheless, FACS analysis showed that MGT#1 (first generation) and MGT#4 (LSD) are similarly responsive to TNF-alpha, a driver of mesenchymal commitment in GBM cells and of MGT#1-2 activation^14^ (**Fig. 2b**). Of note, MGT#1 exhibits a comparable expression regardless of whether genome engineering relies on lentiviral- or piggyback-based systems (**Fig. 2b**). This indicates that sLCRs’ activity is mainly directed by the synthetic *cis-*regulatory DNA and largely independent of the vector employed.

**Figure 2.**
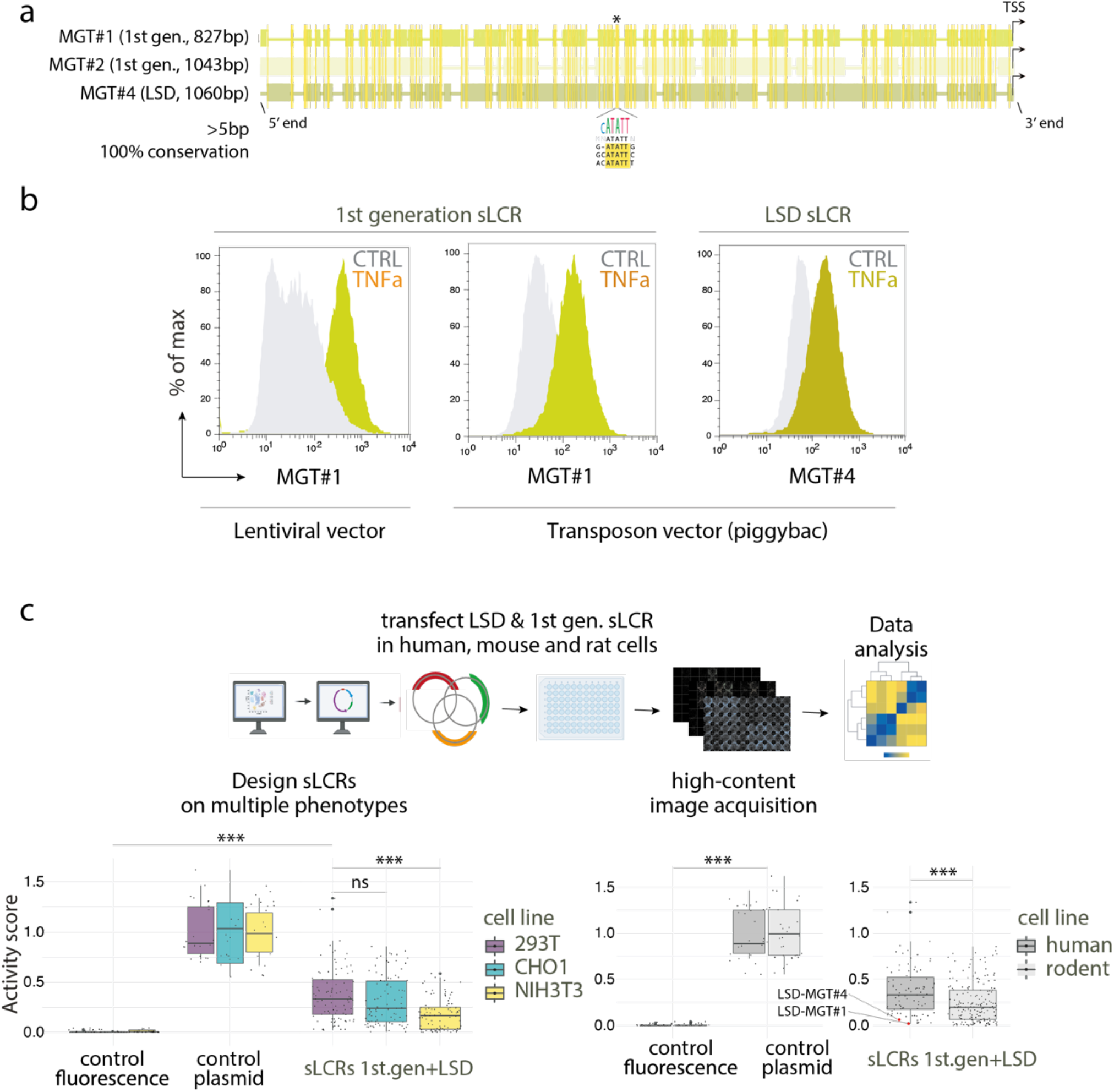
LSD allows designing functional and specific sLCRs. a) Multiple sequence alignment (see Methods) of the 1st generation MGT#1 and MGT#2 and the LSD MGT#4 reporters. The conserved positional overlap is denoted by the asterisk and graphically represented by the sequence logo. b) FACS analysis of MGT#1 (left & center) and MGT#4 (right) sLCRs expression in human glioblastoma-initiating cells with or without TNFa stimulation. Note the similar induction between lentiviral- and transposon-engineered cells for 1st gen. MGT#1 sLCR, and between MGT#1 and LSD-designed MGT#4. c) Above, schematic depiction of the systematic screening of sLCRs designed on diverse phenotypic signatures in three difference species. Lower left, box plot of indicated sLCRs (n=28) transfected in human epithelioid 293T, hamster epithelioid CHO-K1 and mouse fibroblastoid NIH3T3 cell lines. The X-axis shows fluorescence normalized by controls and transfection efficiency per cell line. Each sLCR measurement was assessed in technical replica (n=3). Left, positive (n=5) and negative controls (n=3) denote CFP, GFP, mCherry and iRFP670 expression driven by non-sLCR promoters and fluorescence background in each channel, respectively. Lower right, box plot shows relative activity of human sLCR transfected in human (293T) or non-human (CHO-K1, NIH3T3) cells. Statistics: 2-way ANOVA, followed by Dunnet post-hoc test.

Transcriptional reporters typically consist of candidate enhancers upstream of a minimal promoter^11^. The use of non-specific promoters limits the phenotypic specificity of the reporter. In contrast, no minimal promoters were required to design functional and specific MES GBM sLCRs. To extend this observation, we next used LSD to systematically design sLCRs for a wide range of cell states and pathways’ activation. These include the proneural and classical glioblastoma expression subtypes, an astrocyte-like glioblastoma cell state, ER-stress response, senescence, T-cell exhaustion, disease-associated microglia activation, and epithelial cells’ response to SARS-2 infection. Consequently, the source of signature genes, transcription factors, the choice of reporter genes, an LSD-sLCR-independent selection marker, and vector backbones differed according to the intended outcome. As a result, we generated a broad pool of sLCRs whose connecting thread is being designed by LSD (**Supplementary Table s1** and **Methods**). With such a diverse range of sLCRs, we were able to determine if LSD systematically generates functional reporters. Thus, we synthesized LSD-sLCRs and episomally tested their expression side-by-side with the experimentally validated first-generation reporters in three different cell lines. We transfected 28 different sLCRs into human epithelial HEK293T, mouse mesenchymal NIH3T3, and Chinese hamster ovary (CHO-1) epithelial cells to cover a minimal set of variables that allow assessing transcriptional competence and specificity, including developmental stage, tissue-specificity, and species-specificity. Despite the use of phenotype-agnostic cell types, upon transfection, the median expression of the sLCRs was significantly distinct from the background (**Fig. 2c**). There was no obvious bias associated with the algorithm used to design them or the target phenotype, but the sLCR showed distinct patterns of expression in the three cell types. Overall, reporters designed with the human genome as reference displayed a mild but statistically higher expression in human cells, if compared to mouse and hamster cell lines (**Fig. 2c**). Some LSD-sLCRs were marginally transcribed despite high transfection efficiencies as gauged by the expression independent fluorophore in all the tested lines (**Supplementary Fig. s2c**). This could be interpreted as either a measure of on-target specificity or the lack of transcriptional competence. As exemplified by the case of our mesenchymal GBM sLCRs, those reporters were highly induced by TNF-alpha in GBM cells confirming that they are functional, but lowly expressed in transient transfection of non-glioblastoma cells, supporting their specificity (**Fig. 2b and S2c**). Interestingly, one sLCR scoring very high on-target activity such as PNGT#2 in proneural human glioblastoma initiating cells, was well expressed in human 293T cells but significantly less expressed in non-human cells (**Fig. 2c**), suggesting that the species discordance might affect the output reporter activity.

Thus, LSD generates reporters whose specificity is linked to the source of signature genes, transcription factors and cell model system, and whose transcriptional competence is independent of minimal non-specific promoters.

### Benchmarking LSD by *cis-*regulatory score ranking towards defining basic principles of sLCR design

Given the ability of the LSD approach to optimize TFBS complexity within selected CREs, we next sought to exploit glioblastoma first-generation validated MGT#1-2 reporters^14^ and proneural-to-mesenchymal transition (PMT) as a benchmark to predict the functionality and specificity of newly LSD-designed sLCRs.

First, we set out to estimate an indicative number of distinct TFs that should be represented by cognate TFBS in a given sLCR, in order for it to be functional and specific. To this end, we set out to determine background TFBS complexity by using different randomization strategies. First, we sampled TFs from the overall pool of database-annotated human TFs, while maintaining the signature genes constant (termed Random TF; **Fig. 3a**). Second, we randomly selected matched-sized sets of input genes from the human genome (GRCh37/hg19) and maintained the MGT#1-2 TF list (termed Random Sig.-TF; **Fig. 3a**). We then calculated, for each of the two scenarios, and for increasing numbers of sampled TFs, a measure of specificity, defined as the fraction of TF-genes included in the designed sLCR, and that we annotated as MGT factors. This fraction increased as a function of the input number of random TFs, and this trend was more marked when the MGT#1-2-4 signature genes were used (**Fig. 3a**). Importantly, all mesenchymal sLCRs (MGT#1-2-4), which were designed with less than one hundred TFBS, covered >50% of the entire TF repertoire, and the observed/expected ratio was superior to both the background and the phenotypically distinct classical and proneural reporters (**Fig. 3a**). Interestingly, despite the fact that the MGT#4 reporter was designed by LSD on a different TFBS list, it outperformed the first generation MGT#1-2 on their specific TFBS input list (**Fig. 3a**). This was never the case for other LSD-designed sLCRs, PNGT#3 and CLGT#3 (**Fig.s3a-c**). In fact, using their specific signature and TF-gene inputs, they all showed an observed/expected TFBS ratio above both background and other functional reporters designed to have specificity for a different phenotype (**Fig. s3a**). Hence, the LSD-sLCR approach outperformed the supervised selection by enriching the number of TFBS detected above the signal-to-noise ratio, even at large TF numbers.

**Figure 3.**
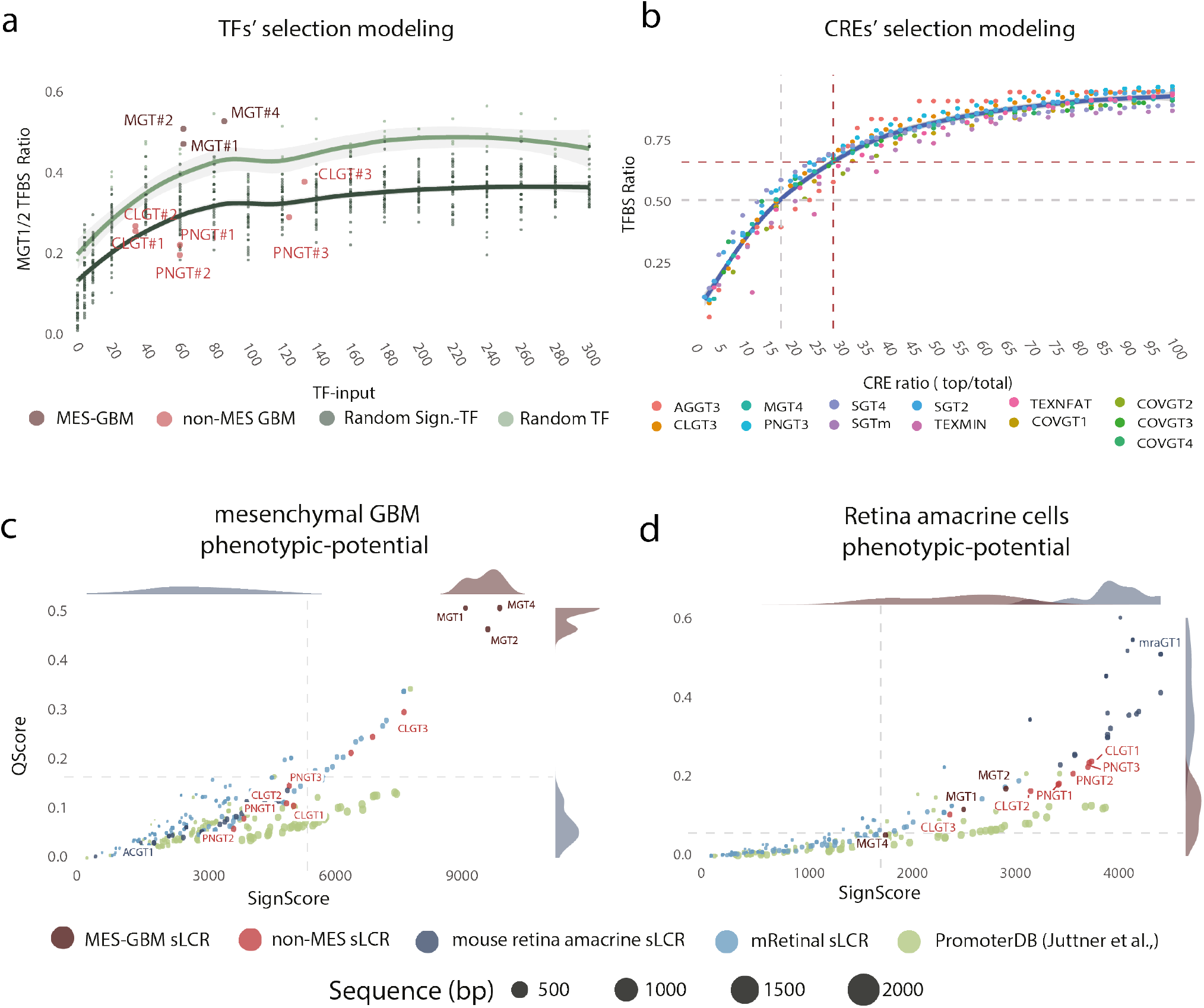
Towards defining endogenous and synthetic reporters’ phenotypic potential via TFBS enrichment ranking. a) Scatter plot showing the mesenchymal sLCRs TFBS affinity ratio for on-target, off-target and scrambled sLCRs. The Y-axis indicates the observed/expected ratio (i.e., MGT#1-2 observed/input TFBS). The X-axis denotes the number of input TF. First-generation and LSD-sLCR are indicated. Scrambled sLCR were designed using LSD and input from random sampling of TFs from the general pool of annotated human TFs (random TF) or random selection of genes from the human genome (random Sign-TF). Fitted lines indicate LOESS regression. b) Scatter plot showing the TFBS affinity ratio as a function of increasing numbers of CREs. Values are calculated for each functional sLCR assessed experimentally (Fig. 2). Logarithmic regression was used to fit the curve. The grey dashed line indicates that the CRE ratio is >50% of TFBS with R2 = 0.96 and the coloured line marks MGT#4. c) Scatter plot showing the signature score (x-axis) and affinity score (y-axis; see Methods) of the indicated reporters for the mesenchymal phenotype. Note the antagonistic phenotypic scoring of glioblastoma and neural retina amacrine cell reporters. d) Phenotypic scoring of the same reporters in (c) for a retina amacrine cell phenotype.

Beyond the threshold of one hundred TFs, our randomization approach suggests that while the number of target TFBS proportionally increases with the size of the sLCR, this does not affect the fraction of specific TFs recovered (**Fig. 3a**). This prediction is compatible with the idea that unnecessary long sLCRs are prone to unpredictable regulatory activities, suggesting that the length of an sLCR is a critical determinant of its specificity and sensitivity. Thus, we next set out to define a minimal number of individual CREs that would predict a functional sLCR while aiming at maximal TFBS potential. To this end, we quantified the marginal information gain (i.e., the number of distinct TFs included in the sLCR, as quantified by the cognate TFBS) of increasing the number of individual CREs to be included into an sLCR. This analysis integrated a total of twenty-eight functional sLCRs targeting various phenotypes (**Fig. 2d**; **Supplementary Table s1**). Fitting a model to the relationship between the TFBS fraction and the number of CREs included using LASSO (R > 0.9; **Fig. 3b**), retrospectively determined that the top 20% of output CREs is sufficient to represent >50% of the maximal regulatory potential in a given phenotype. For example, for the herein validated MGT#4, we selected 7 out of 24 CREs (29%), for a total length of the sLCR of 1060bp, and an observed/expected ratio of 0.56. Taken together, the results of this analysis suggest that, if input sets with comparable sizes are used for novel sLCR design, merging 25-30% of the top-ranking CREs may maximize the chances of obtaining a functional sLCR, with minimal size and thus ectopic activities (see discussion).

Finally, we tested whether the validated reporters could inform a model to predict the phenotypic specificity of a given candidate *cis-*regulatory DNA sequence, endogenous or synthetic. To this end, we established a *cis-*regulatory score ranking. Such ranking is based on the correlation between a score summarizing the overall affinity of each sLCR to the phenotype-specific TFs (termed Qscore), and a score proportional to the phenotype-specific expression of the genes in the corresponding input signature included in the final design of each sLCR (termed SignScore; see **Methods**).

Using the TFs and input gene signatures employed to design the MGT#1-2 reporters (intended as a validated proxy of the mesenchymal GBM phenotype), the sLCR based on the mesenchymal TFs and signature (including MGT#4) outranked all the remaining reporters (**Fig. 3c**). To test the specificity of this ranking strategy, we introduced reporters potentially marking a phenotype distant from those of GBM-subtypes. We used LSD to design sLCRs to map amacrine cells starting from murine single-cell RNA-seq profiling^19^ and compared these reporters to a set of validated synthetic reporters generated by various approaches to perform gene therapy in mouse retinal cell types^69^. Importantly, mouse retinal reporters outranked glioblastoma sLCRs in the amacrine phenotype ranking, while they sat at the bottom of the mesenchymal glioblastoma phenotype ranking (**Fig. 3c-d**). Likewise, when this analysis targeted classical or proneural GBM TFBS selections, their respective reporters outperformed all the others (**Fig. s3b**).

Overall, by using our validated sLCRs as a benchmark, we set up a series of computational strategies that can aid in the design of functional reporters and measure the *cis-*regulatory potential affinity of synthetic and endogenous reporters to their target phenotype.

### LSD incorporates single-cell RNA-seq as signature gene input for LSD

Having established the empirical performance of LSD on bulk RNA-seq and reference genomes, we sought to exploit scRNA-seq data as signature gene inputs for LSD.

Since glioblastoma subtypes were recently reassessed as distinct cell states using single-cell RNA-seq^20^, we next used these meta signatures to generate sLCRs from scRNA-seq inputs. As a resource for glioblastoma-specific TFBS, we resorted to either bulk or scRNA-seq lists and designed the reporters for the four scRNA-seq states.

Whereas the mesenchymal glioblastoma subtypes and states are substantially overlapping, and ssGSEA analysis indicated that MGT#1-2 (1^st^ gen) and MGT#4 (LSD) are already representative of this state (**Fig. s3d** and ^14^), the relationship between non-mesenchymal glioblastoma subtypes and states is unclear. Computationally, proneural and classical sLCRs broadly span through two states, but the AC-like sLCRs clearly represent the classical GBM subtype (**Fig. s3e** and ^14^). Hence, the resource provided herein (**Supplementary Table s1**), may be helpful to define critical cell intrinsic signaling and cell fate changes upon perturbation, which may be particularly useful to study the significance of specific cell states.

### LSD integrates 3D contact maps and DNA accessible in chromatin as custom input

Chromatin accessibility is a primary determinant of cell type-specific *cis-*regulatory activity, and accessible TFBS are more likely to bind cognate *trans*-regulatory factors^21,22^. The 3D genome organization guides the spatiotemporal function of enhancers in the mammalian genome^17,18^. The increasing availability of cell type-, developmental state- and disease-specific chromatin structure maps prompted us to test whether sLCRs may be designed with the aid of such input datasets.

First, we designed the mesenchymal glioblastoma MGT#4 LSD-sLCRs from four alternative input combinations (**Fig. 4a**). The nearest-neighbor CTCF binding sites approach applied to the full extent of the reference genome is the standard approach described above (**Fig.4a, I**). MGT#4 design iterations were obtained by either restricting the reference genome to the cancer-specific accessible genome, as defined by ATAC-seq profiles^23^ (**Fig. 4a**, III and IV), or by extending the space for candidate cis-regulatory domains to tissue-specific TADs ^24^ (**Fig. 4a**, II and IV). We used four indpendenent sources of ATAC-seq data, including mouse regulatory regions^25^. The signature genes and TFBS input were identical to those used in MGT#4 LSD-sLCR. Therefore, this analysis generates a set of distinct LSD-sLCR designs potentially targeting the same phenotype.

**Figure 4.**
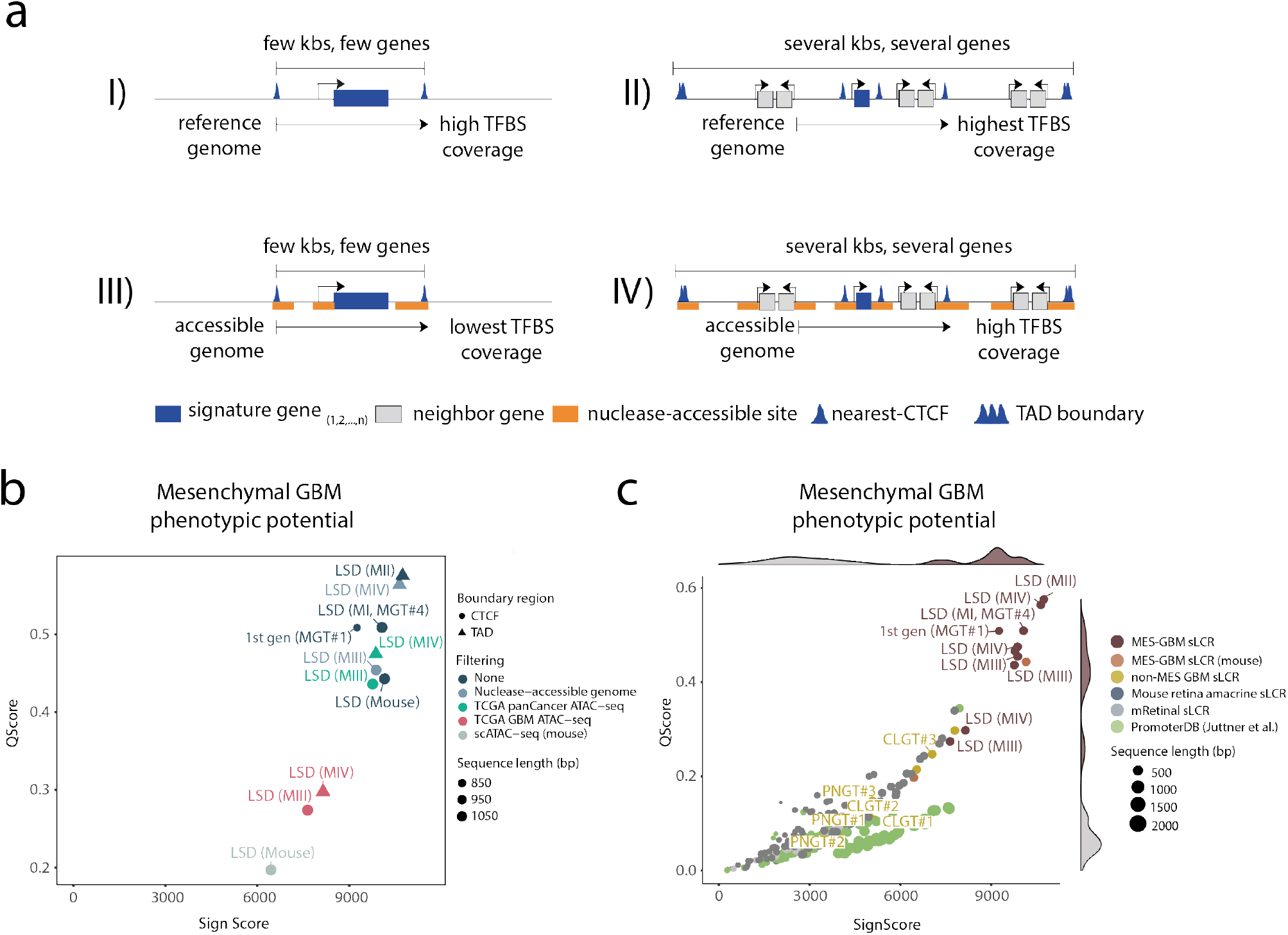
Integrating chromatin accessibility and 3D contact maps as input for LSD. a) Schematic representation of alternative LSD input combination models. b) Scatter plot showing the signature score (x-axis) and affinity score (y-axis; see Methods) of the indicated reporters for the mesenchymal phenotype. Note the improved on-target score for a mesenchymal sLCR designed by LSD using model II (i.e. MGT#5). c) Scatter and density plots of data in (b) with the addition of non-specific phenotypic reporters. Note that the mesenchymal phenotypic space is occupied by most mesenchymal reporters and that the including of accessibility and 3D contact data marginally increased or decreased sLCR scoring.

To compare the sLCRs designed by LSD according to the four different models (**Fig. 4a**), we constrained sLCR size to that of the validated MGT#4 reference. We computed the cis-regulatory score affinity to the MGT#4 target phenotype for all the mesenchymal GBM sLCR designed with the above input iterations to all the available reporters. *Cis*-regulatory score ranking shows that applying interactions to the input DNA changes the *in silico* specificity of the reporters (**Fig. 4b**). Yet, all mesenchymal reporters occupy a distinct space from phenotypically distant sLCRs and background sequences, and the mesenchymal sLCR designed with the largest input for TFBS search slightly outperformed validated reporters using the standard inputs (**Fig. 4c**).

To complement the *cis*-regulatory ranking with a statistical approach that estimates data distributions with limited sampling and in absence of major assumptions, we next used a bootstrapping analysis strategy^26^. We included all the different MGT#4 iterations, non-mesenchymal sLCRs and phylogeny-distant DNA control regions from the SARS-CoV-2 viral genome. We conducted iterative random sampling of MGT#4 TFBS enrichment at each LSD reporter under testing (n=1,000) using a fixed TFBS number (25% of the total, 231, with replacement). At each iteration, the total value of the random TFBS selection was calculated, thereby creating a distribution of TFBS enrichments for each LSD reporter. Then, each data distribution was compared against the others for each LSD reporter (greater=TRUE). The bootstrapping analysis established a hierarchy of significant pairwise correlations between distributions, in descending order. The significance directly correlated with the size of the input reference for TFBS search (**Fig. s4b-c**). The number of significant events occurring decreased when the size of the ATAC-seq data was smaller than the reference genome and improved when the size was comparable (e.g. MIII; **Fig. 4b-c and s4d**). In other words, constraining the TFBS search to a limited ATAC-seq dataset results in fewer significant interactions than when using the unrestricted reference genome. This can be interpreted as the limited likelihood of covering the entire TFBS repertoire in a short sLCR, which increases the number of CREs required to represent the target cis-regulatory potential (i.e., the overall size of the sLCR). Consistently, when LSD was restricted to the GBM ATAC-seq genome and the nearest-neighbor CTCF boundaries, the optimal sLCR designed by LSD outcome should be ∼16% larger than MGT#4. The use of larger *cis-*regulatory pools, such as the pancancer ATAC-seq or a curated set of nuclease-accessible genome regions from various ATAC-seq and DNAseI-seq maps, relieves such a constraint. In fact, the combination of TADs and the pancancer ATAC-seq datasets ranks similarly or slightly outperforms MGT#4 *cis-*regulatory potential, as gauged by both cis-regulatory score ranking and bootstrapping analyses (**Fig. 4b-c** and **s4b-d**).

In summary, provided that the size of the custom cis-regulatory datasets for TFBS search is comparable to the unrestricted reference genome, LSD well integrates 3D contact maps and DNA accessible in chromatin inputs toward extending the use of synthetic genetic tracing to the validation of functional experimental datasets.

### LSD enables prioritizing cell-state-specific drivers in combination with genome-wide gain-of-function CRISPR screens

The generation of cell state-specific reporters enables the discovery of cell state regulators by hypothesis-driven approaches and also by unbiased genetic screens^14^. To illustrate the experimental opportunities enabled by designing sLCRs to represent multiple cell states, we performed genome-wide pooled CRISPR activation screens (CRISPRa) in Isocitrate dehydrogenase wildtype human glioma-initiating cells (IDH-wt-hGICs) with either MGT#1, MGT#4 or PNGT#3 as a readout. This strategy potentially allows the discovery of genes whose transcriptional amplification leads to a mesenchymal or proneural cell fate commitment.

A CRISPRa library targeting 18,915 human genes via 104,540 on-target and 1,895 control gRNAs (∼5 gRNAs/gene)^27^ enabled us to carry out a genome-scale gain-of-function phenotypic screen. The presence of a fluorescent marker in each vector directly supports that the conditions of transduction respected the one guide per cell rule (multiplicity of infection[MOI] <0.5), while maintaining a minimal library representation throughout the cell culture experiment (i.e. ∼200x). We FACS isolated the cells with the highest and lowest expression of the reporter for each GBM subtype sLCR after seven passages and fifteen days of gene activation through specific gRNAs, a time frame designed to permit transcriptional activation and cellular reprogramming (**Fig. 5a**).

**Figure 5.**
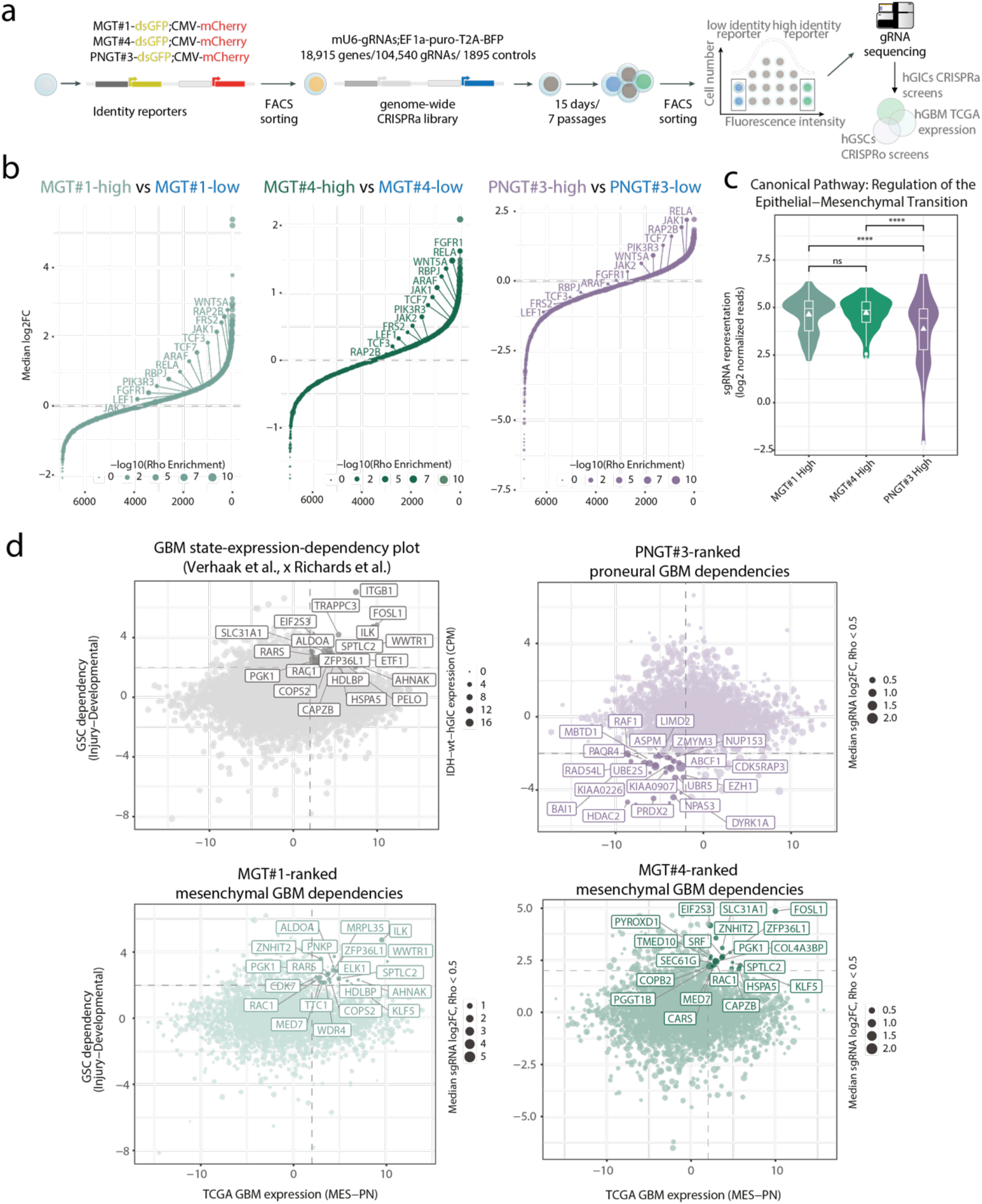
Discovery of cell-state-specific drivers by convergence of LSD, genome-wide CRISPR activation and expression and essentiality maps in patients. a) Workflow of the CRISPR activation screens. b) Rank plot showing median log2-fold changes for gRNAs between the indicated sLCR-high and -low fractions. Labels highlight genes defined as connected to EMT terms by Ingenuity Pathway Analysis. The analysis was limited to genes expressed in target cells (CPM>5). c) Violin plot of the mean gRNA read-count distribution for EMT-related targets highlighted in (b). P-values denote significance of pair-wise comparison (Wilcox.test, padj<0.05). d) Scatter plot showing GBM subtype-specific expression (X-axis) and GBM stem cells dependency (Y-axis). Dot size represents either gene expression in naive IDH-wt-hGICs (top left), or median log2-fold change of the CRISPRa targets (all other plots). For each sLCR, highlighted genes represent candidate GBM subtype-specific biomarkers, dependencies and drivers. For candidate drivers of mesenchymal GBM overlapping between MGT#1 and MGT#4 screens see Fig. s3.

At the experimental endpoint, there was a substantial equilibrium between guide-bearing and non-infected cells and the reporter expression appeared independent of the library (**Fig.s5a**). This suggests that the library overexpression might have introduced mild non-autonomous cell fate changes but – more importantly – it excluded bleed-through technical artefacts that may complicate the FACS sorting of phenotypically different cells based on fluorescence. Next, we subjected the candidate outliers to FACS sorting, genomic DNA extraction, PCR amplification and sequencing of the gRNA library, similar to our recent sLCR-driven phenotypic loss-of-function screens ^14,28^. Sequencing of the plasmid library confirmed that the qualitative representation was maintained. Quantitatively, each condition clustered independently, indicating that sorting subsets of cells by reporter expression introduced a selective segregation of the gRNAs (**Fig.s5b-c**). When comparing all cell-based conditions to the library, only a limited subset of genes scored as significantly differentially enriched There was a notable enrichment of cell cycle promoters in cells, including Cyclin D1 and Cyclin D3 (**Fig. s5d**), indicating that active gRNAs in the library can functionally amplify biological programs.

Unbiased analysis of the dataset using robust rank aggregation and enrichment over the reporter-low fraction, uncovered genes whose potential amplification by multiple gRNAs, could drive reporter-specific upregulation (**Fig. 5b, s5f and Supplementary Table s2**). Strikingly, we found RELA among the MGT#4 screen top hits (**Fig. 5b, s5f and Supplementary Table s2**): RELA is one of the NF-kB transcription factors that we previously identified as an MGT#1 regulator under homeostatic conditions in glioblastoma-initiating cells in both focused loss-of-function experiments and genome-scale CRISPR KO screens^14^. Since we previously showed that MGT#1 is inducible via multiple signaling pathways in *in vitro* settings ^14^, we next focused our analysis on common MGT#1 and MGT#4 candidate hits as a means to globally illustrate the discovery power of the combined LSD-CRISRPa approach. Ingenuity Pathway Analysis (IPA) revealed that the mTOR (padj=1.44544E-07), BEX (padj=3.39E-06), WNT (padj=2.45E-04) and EMT canonical (padj=6.17E-06) pathways connected genes defined by two gRNAs with a significant positive enrichment in both MGT#1 and MGT#4 screens. In particular, the the EMT pathway gene set was significantly more enriched in MGT#1 and MGT#4 screens when compared to PNGT#3 (**Fig. 5b-c**). This suggests that reporter-expressing cells correctly captured the gain-of-function activity imparted by CRISPRa gRNAs that are likely candidates for a PMT.

To overcome the limitations imposed by each experimental platform, we reasoned that MGT#1-4 and PNGT#3 sLCR drivers which converge on differential expression in glioblastoma biopsies (TCGA GBM, Verhaak et al., ^29^) and glioblastoma stem cell state-specific dependencies identified by systematic CRISPR genome-wide KO screens^30^, would identify genuine phenotypic drivers (**Fig. 5a**). Indeed, constraining the analysis to the GBM-subtype-specific dependencies leads to a 156.32 and 171.95 fold enrichment over expectations for overlap with MGT#1 and MGT#4, respectively (hypergeometric p-value = 1.36E-16 and 5.39E-17, respectively; **Fig. s5g**). Ranking of the top 500 hits by fold changes, maintained a significant overlap between MGT#1 and MGT#4, albeit far less enriched (7.8%, hypergeometric p-value = 1.77E-09). Together, these analyses indicate that convergent hits from these orthogonal experiments effectively restrain the background contributed by the use of different disease models and technical limitations associated with either experimental approach (see discussion). Importantly, both screens featured the eight shared novel mesenchymal candidate drivers (**Fig. 5d, s5g**), and each reporter identified specific candidates, including the extensively validated WWTR1^31^ by MGT#1, and FOSL1^32^ by MGT#4. Both WWTR1 and FOSL1 are well established mesenchymal-GBM transcription factors and could have been prioritized based on the intersection of the Verhaak and Richards datasets and prior knowledge. Instead, the other hits share features that make them likely but non-obvious novel candidates for a cell-intrinsic PMT driver function. RAC1 was previously shown to connect mTOR-dependent growth control and STAT3 signaling in NF1-deficient cells^33^. PGK1 is a well-established HIF1a-regulated gene, and its upregulation is more obvious in non-stem glioma cells under hypoxia as compared with matched GSCs^34^. The zinc-finger protein ZNHIT2 is instead poorly characterized, and appears to be involved in the spliceosome machinery, and potentially connected to mTOR-regulated response to energy- and nutrient-sensing stress response^35^. Overall, among the eight shared hits featured by our orthogonal analysis as drivers of a mesenchymal program activation, 37.5% (3 out of 8; KLF5, MED7, ZFP36L1) point to transcription and 62.5% (5 out of 8; PGK1, RAC1, COBP2, ZNHIT2, SPTLC2) point to regulation of metabolism, which is an inseparable feature of the mesenchymal program, as also recently discovered in a pathway-based GBM subtype classification ^36^.

The intersection of the Verhaak and Richards datasets alone does not appear to define obvious *bona fide* proneural regulators (**Fig. 5d**). Importantly, the orthogonal combination of all the three approaches defines a subset of thirty-eight hits. Globally, by Ingenuity Pathway Analysis, these hits featured the activation of the EIF2 pathway (p=5,64E-09) microRNA biogenesis (p=7,93E-08) and Hugntington’s disease signaling (5.73E-07) pathways, and the MYC transcriptional program activation (p=5.3E-15). The high proneural identity displayed by our *in vitro* models might limit the discovery power of the PNGT#3 screen and suggests that mesenchymal GSCs or signaling may provide better ground for screening proneural amplifiers. However, the identification of the adhesion G protein-coupled receptor B1 (ADGBR1) also known as brain-specific angiogenesis inhibitor 1 (BAI1), a synaptic receptor whose overexpression limits neo-vascularization and xenograft formation ^37^ and correlates with APLRN-driven invasive potential in proneural glioblastoma cells ^38^. Hence, BAI1 known function is consistent with a role in proneural differentiation.

Together, the experimental combination of LSD-designed sLCRs, CRISPR screens and with patients’ molecular maps uncovered the *bona fide* regulators of cell states that connect the tumor genotype to its molecular phenotype. This showcases the utility of LSD as a generally applicable framework to study cell identities and cell state changes.

## Discussion

Here, we present LSD as a framework to streamline the tracing of cell identities and state changes for complex phenotypes by synthetic genetics. LSD scans the reference genome for candidate *cis-*regulatory DNA using one list of biomarkers and one of TFs defined as active in the cell type or state of interest. Using previously validated GBM sLCRs^14^ as benchmark, it is shown that the LSD primary output outperforms previous designs *in silico*, and is functional and specific in head-to-head experimental validation, including a genome-wide CRISPRa screen. The seamless generation of genetic tracing reporters for distinct cellular phenotypes by LSD represents a resource for future experimental validation (**Supplementary Table s1**). Our modelling approach on validated mesenchymal GBM sLCR shows that both high levels of target TFBS (Qscore) and enrichment of TFBS at signature genes (SignScore) are higher in mesenchymal LSD reporters than in those specific toward different phenotypes or previous selection. This underscores the importance of curated TFBS inputs to advance specificity of the reporters, which will likely benefit from the recent development of CUT&RUN/TAG for TF footprinting ^39^. Broadly, despite the LSD ability to generate a selection of CREs that outperforms random selections, a better understanding of sequence-specific gene regulation and a high-quality input are likely critical for effective learning and ranking by LSD. Hence, iterative refinement that combines LSD upstream computational design tools^40^ and downstream massively parallel testing^4,12^ holds the potential to improve synthetic reporter design even further.

The advent of high-throughput chromatin accessibility profiling enabled mapping gene regulatory landscapes in healthy and diseased cell states but outpaced the development of experimental methods to directly test hypotheses generated from these data. Moreover, *bona fide* cell states identified by single-cell data *ex vivo* require *in vivo* validation, as artefactual signatures may be confounding ^41^. Previous methods successfully leveraged *cis-*regulatory DNA and expression datasets to deploy cell-type-specific enhancers and enabled genetic tracing and perturbation of gene function across mammalian cell types^9,11^. Endogenous or minimal viral promoters were key to generating functional and specific synthetic promoters in past approaches ^11^. Similar to our previous design of sLCRs ^14^, LSD generates functional reporters without minimal promoters, which is a key advantage to increasing phenotype specificity. Moreover, we developed a ranking system to assess the on-target potential of genetic reporters. With this combination, we designed the second generation of glioblastoma sLCRs, as well as other sLCRs targeting a diverse set of molecular phenotypes. Supporting this approach as broadly applicable, one such sLCR designed to address the molecular response of epithelial cells infected by the SARS-CoV-2 recapitulated the basic principle of epithelial cells’ innate immune responses and uncovered expected and unexpected pharmacological regulators of such response (Jiang B., Schmitt M., et al., *manuscript in preparation*).

Whether the use of custom inputs will serve validation purposes or is key to improving the functionality and specificity of the sLCRs will require systematic, large-scale testing. Nevertheless, our focused statistical approach indicates that a large dataset of validated *cis-*regulatory DNA is compatible with focusing the discovery to regulatory elements that may preserve evolutionary-shaped regulatory logic without assuming that such logic plays a determining role in *cis-*regulation by synthetic DNA. Likewise, whether the use of an input non-species-matched genome will enable synthetic genetic tracing development and omics validation in model organisms, such as when we used mouse ATAC-seq data to design a mesenchymal GBM sLCRs, warrants experimental testing. Nevertheless, incorporating experimental inputs – such as chromatin accessibility and 3D contact maps – showcases LSD’s versatility and aids in defining a better on-target CRE selection.

We previously showed that sLCRs connect changes in the mesenchymal state of glioblastoma or lung cancer cells to the loss of function caused by CRISPRout or CRISPRi in pooled screens ^14,28^. Here, we expanded the use of sLCRs in phenotypic pooled screens to gain-of-function by CRISPRa. Unlike the proneural sLCR, the mesenchymal sLCRs promoted the representation of gRNAs targeting EMT genes among dCas9-VPR and gRNA-bearing cells. In combination with patients’ gene expression profiles ^29^ and functional screens in patient-derived cells ^30^, genome-scale CRISPR activation in sLCR-containing cells pointed to novel candidate drivers of individual glioblastoma transcriptional phenotypes, including genes not previously described as drivers. This illustrates the utility of combining descriptive and functional information to prioritize candidate targets and biomarkers for validation. Of note, this approach may be particularly helpful in understanding the molecular foundation of glioblastoma subtypes and states whose significance is still unclear. In line with the reclassification of glioblastoma states by pathway activation^36^, our analysis uncovered several metabolic regulators. The identification of known mesenchymal GBM drivers such as WWTR1^31^ and FOSL1^32^ by MGT#1 and MGT#4, respectively, denotes the specificity of the sLCRs and the need for improving the signal-to-noise ratio in phenotypic pooled screens. The constant development of powerful dCas9 effectors and of arrayed gRNA libraries is likely to improve the discovery power of genome-wide phenotypic screens ^42^. Arrayed screens are also anticipated to reduce the influence of non-autonomous phenotypic changes in pooled screens, which are responsible for neutral gRNA background recovery in multicellular 3D models like the one shown here.

Applying Boolean logic gate strategies^43^, including regulators of mRNA translation and stability as well as protein homeostasis regulation, will further enhance accuracy and specificity of complex phenotypes’ synthetic genetic tracing.

In conclusion, we expect LSD to provide a simple and scalable approach to perform genetic tracing in complex multifactorial settings, enable validation experiments of omics approaches on equal complexity terms, as well as to study fundamental questions underlying transcriptional regulation.

## Acknowledgments

We are grateful to L. Li, H. Naumann, M. Grossman and the MDC FACS and genomics technology platform for technical support and A. Akalin and S. Mzoughi for critical reading of the manuscript. Data analyses include data generated by the TCGA Research Network: https://www.cancer.gov/tcga. CC, MJS and YD are graduate students with Humboldt University, BJ and SK with Charitè Medical University. SK and YD are affiliated with the Berlin School of Integrative Oncology (BSIO) at Charitè. The GG lab acknowledges funding from MDC, Helmholtz (VH-NG-1153) and ERC (714922).

## Methods

### Datasets

TCGA data were from dbGaP database of Genotypes and Phenotypes (dbGaP) accession phs000178. The GSCs dependencies were from the supplementary material in Richards et al. CRISPRa screen data generated in this study are attached in Table S2.

### LSD algorithm

The LSD algorithm takes a list of position weight matrices (PWMs) along with the corresponding cognate TF-genes, a list of marker genes of a target phenotype, and the reference genome of the organism of interest, and it generates a list of naturally occurring, putative *cis-*regulatory elements, that are then used to assemble the synthetic-reporter for the target phenotype. The algorithm can be divided into three steps. In step I, LSD generates a pool of potential CREs with a fixed length within user-defined regulatory landscapes (default is a 150bp window sliding with a 50bp step). In step II, LSD assigns TF-binding sites to the CRE pool using FIMO (default *--output-pthresh 1e-4 --no-qvalue*), and creates a matrix of putative CREs x TFBS. In step III, LSD ranks and selects the minimal number of CREs representing the complete set of TFBS. For that purpose, it iteratively sorts and selects the best CRE based on the overall TFBS affinity and diversity among the input TFs showing high affinity for the CRE. Starting from the ranked CREs, LSD selects the highest-ranking CRE defined by the sum of the affinity score (-log10(p-value)) and TFBS diversity (number of distinct TFs represented in the predicted TFBS). Subsequently, it removes the selected CRE and the corresponding TFBS from the CRE x TFBS matrix and repeats the selection. This continues until either none of the CRE or of the TFBS is left. In the ranking, priority is given to CREs proximal to known TSS to increase the chances of successful transcriptional firing using the same strategy as above. TSS were defined based on FANTOM CAGE peak BED files (RRID:SCR_002678; Human: http://fantom.gsc.riken.jp/5/datafiles/phase1.3/extra/TSS_classifier/TSS_human.bed.gz and TSS-containing CREs were defined based on overlap with genome-wide pooled CRISPRi libraries^44^. Finally, LSD returns an ordered list of the selected CREs, together with a representation of the TFBS scores (**Fig. 1A**). The framework to run the LSD algorithm is available at: https://gitlab.com/gargiulo_lab/sLCR_selection_framework.

### LSD reporters design

Reporters were designed using the LSD method are indicated in supplementary Table S1. First-generation GBM-sLCRs^14^ were designed by manual integration of a selection of top-ranked CRE. LSD GBM-sLCRs (**Fig. 1**) used as input the first generation sLCR gene signatures, and a selection of subtype-specific TFs (high-expressed TF genes, > quantile 75%; https://meme-suite.org/meme/) obtained from TCGA-GBM patients’ RNA-seq expression profiles (RPKM-UQ, TCGA, phs000178.v3.p3). Random Sign/TF sLCRs were generated by integrating different sizes of randomly selected human genes (three different sampling of 10,20,50 and 100 different genes) and TF genes (four sampling of 1 to 301 different TF). Random TF (only) used MGT#1-4 signature genes to generate the CRE pool. AC-like state LSD-sLCRs were generated using Neftel et al. AC-like signatures^20^ and CLGT#1-2-3 TFBS. Retinal LSD reporters were generated using retina-specific cell-population markers genes (GSE81905) TF genes determined at different thresholds (> quantile 60%-88%). hg19 (GBM) and mm10 (retina) assemblies were used to extract genomic positions.

Multiple sequence alignment was conducted with four different tools (MUSCLE, Clustal Omega, T-Coffee and MAFFT in SnapGene) with similar outocomes. MUSCLE output is reported in **Fig. 2a**.

### Phenotypic potential inference by on-target TFBS scoring

The inference of the specific phenotypic potential of each LSD-reporter was defined as the linear correlation between the Qscore (affinity-score) and the SignScore (phenotypic-score). The affinity-score (QScore) was calculated for each reporter as the sum of the TFBS-affinity using FIMO (default *--output-pthresh 1e-4 --no-qvalue*) given a specific set of TFBS (e.g., MGT#1-2 TFBS) and normalized by the sLCR sequence length and the ratio of observed/expected TFBS (∑ (FIMO-score) * sequences length)/ (observed/expected TFBS))). In contrast, the phenotypic score (SignScore) was defined as the ssGSEA-enrichment value calculated on target expression profiles (ssgsea.norm=FALSE) using the signature genes, and normalized by the ratio of observed/expected TFBS (ssGSEA-score * (ratio observed/expected TFBS)). Scatter plots were generated using ggplot2 on an R v.3.6 environment.

### Evaluation of the CRE selection

To model the number of CRE required to account for 50% of the total TFBS potential, we used experimentally validated LSD vectors and fitted a lasso regression model. To begin, we generate a list of all the top-CRE sequences for each LSD-sLCR generated. Then, we used FIMO (default —output-pthresh 1e-4 —no-qvalue) to map their corresponding TFBS to each sequence and calculated the ratio of observed/expected TFBS for each top-CRE selection. Finally, we use the lasso function to fit the top-CRE and TFBS ratios. The lasso function was used for the fitting and the the ggplot2 package to generate plots (R v.3.6).

### Comparison of LSD reporters

The analysis of differences between GBM-subtypes enrichment scores was generated by comparison of ssGSEA values (GSVA v.1.32, ssgsea.norm=TRUE, and default parameters) for each synthetic-reporter. To evaluate the similarities between GBM-sLCR transcriptional potential, we evaluate the correlation between TFBS-affinity of all GBM-TFBS and associated TF/TFBS for each GBM-sLCR. The evaluation of the signal-to-noise ratio in ME-GBM was calculated by defining the MGT#1-2 observed/expected TFBS ratio FIMO (default *--output-pthresh 1e-4 --no-qvalue*) using GBM-sLCR and random sLCR. The evaluation of the correlation between the phenotypic-potential in GBM-ME and Amacrine-cell was generated by computing the phenotypic-potential correlation using MGT#1-2 and Amacrine cell TFBS (exclusive high-expressed TF > 75%; below) on each synthetic reporter. The signature enrichment was calculated by using the average of ME-GBM patients’ expression profiles and Amacrine cells single-cell expression profiles. PromoterDB sequences were retrieved from publication (https://data.fmi.ch/promoterDB/) and integrated into the phenotypic-potential analysis with other design LSD-sLCRs. The inference of the phenotypic potential of LSD ACL reporters was generated by using CLGT#1-2 TFBS and the average of CL-GBM patients’ expression profiles. Graphical representations were generated using ggplot2 on an R v.3.6 environment.

### Integration of random LSD-sLCR

To evaluate the signal-to-noise ratio for the LSD-generated reporters, we generated LSD reporters using randomly selected TF and signature genes. First, we sampled using the sample R function different sizes of TF (from 1 to 301 by 20 each, without replacement) from the same input TF database used to generate the LSD and sLCR reporters (repository). Then, in the same way, we sampled different sizes of signature genes (10,20,50, and 100 genes) using the same assembly to generate the sLCR reporters (hg19).

To generate the reporters, we used the LSD pipeline with default parameters for different combinations of inputs (e.g., random TF and random signature genes). To evaluate the combination, we evaluated the TFBS on the included LSD-sLCR and first-generation sLCR. The model of the distribution was generated using geom_smooth (default parameters). Finally, the graphs were created with ggplot2 in R v.3.6 environment.

### Evaluation of the Glioblastoma states

To evaluate the distribution of ssGSEA score on Glioblastoma single-cell expression profiles, we used IDHwtGBM.processed.SS2.logTPM.txt from Neftel et al.^20^. The evaluation of the score integrates signature and TF gene expression on those cells. We maintained the separation between AC/MES and OPC/NPC to differentiate between states. The ssGSEA score was generated using the GSVA R package. The graphs were created with ggplot2 in an R v.3.6 environment.

### Bootstrap analysis of LSD reporters

We used a bootstrap approach to rank the differences between LSD reporters generated with an identical set of Signature genes and TF but with distinct boundary regions and known-CRE filter conditions (Results, Fig.5a). To begin, we generated several LSD reporters (Methods) using the MGT#4 signature genes and CTCF or TAD-defined boundary regions ^24^. The LSD approach allows for filtering regions of interest by determining the overlap between those regions and the CRE. We used this functionality to filter CRE obtained from the LSD pipeline using pan-Cancer ATAC-seq regions^23^. Input genes and TF from MGT#4 were correlated to the mouse and used those as input to generate the mouse LSD reporters. The regulatory regions for mouse LSD were defined from scATAC-seq. After generating the reporters using MGT#4 input, we used CRE to conform the LSD-reporters. We evaluate the sequences by mapping MGT#4 TFBS to each sequence (FIMO; default —output-pthresh 1e-4 —no-qvalue). We used the affinity values for the boostrap analysis. To compare reporters, we conducted a bootstrap analysis. First, the algorithm generated a reporter-specific distribution of TFBS scores by randomly sampling (n=100) TFBS binding to the reporters (25% of MGT#4 TFBS, 60/231). Second, it compares the difference between reporter-specific distribution using the Wilcox test (alternative=greater). Finally, it ranks the reporters according to the number of significant events obtained (adj. p-value < 0.05).

### Human Glioma-initiating cell line (hGICs)

The IDH WT-hGICs were generated by our lab and will be described elsewhere. Extensive histopathological, molecular and tumorigenic potential characterization has been performed and examples are shared with Editors and Reviewers. Briefly, IDH WT-hGICs were generated by transforming human NPC^45^ (kindly provided by R. Glass, LMU), by means of: pRSPURO-sh-PTEN(#1; kind gift from D. Peeper), pLKO.1-sh-TP53 (TRCN0000003754) and pRS-shNF1. For this line, thorough genetic, transcriptional, and epigenetic characterization has been performed, as well as in vivo tumor formation and phenotypic mimicking ability. In vitro, hGICs were propagated as described^46,47^ with one modification. In addition to with EGF (20 ng/ml; R&D), bFGF (20 ng/ml; R&D), heparin (1 μg/ml; Sigma) and 1% penicillin and streptomycin, PDGF-aa (20 ng/ml; R&D) is also supplemented to RHB-A (Takara). This medium composition will be referred to as RHB-A complete. hGICs were cultured at 37°C in a 5% CO2, 3% O2 and 95% humidity incubator.

### Transfection/Transduction

Transfection and transduction of Lentiviral constructs were previously described in detail^47^. Briefly, 12μg of DNA mix (lentivector FH1-MGT#1-mVenus_PGK-H2B-CFP, pCMV-G, pRSV-REV, pMDLG/pRRE) was incubated with the FuGENE-DMEM/F12 mix for 15min at RT, added to the antibiotic-free medium covering the 293T cells, and the first-tap of viral supernatant was collected at 40h after transfection. Titer was assessed using Lenti-X p24 Rapid Titer Kit (Takara) according to the manufacturer’s instructions. We applied viral particles to target cells in the appropriate complete medium supplemented with 2.5μg/ml protamine sulfate. After 12–14h of incubation with the viral supernatant, the medium was refreshed with the appropriate complete medium.

For PiggyBac-vector delivery, AMAXA™ 4D-Nucleofector™ (Lonza, Cologne GmbH) was used for the nucleofection after optimisation of nucleofection conditions (Nucleofector® programs and solutions) for the specific cell line. For each transfection reaction, 1.5μg of pPB[Exp]-mCherry-{MGT#1}>d2EGFP or pPB[Exp]-mCherry-{MGT#4}>d2EGFP, 0.5μg of Super PiggyBac Transposase (System Biosciences, PB210PA-1-SBI) and 0.5μg of piRFP670-N1 (addgene #45457) was used and a final mastermix of DNA and B1.1 buffer with supplements (6mM KCl, 15mM MgCl2, 120mM Na/H2/PO4 pH 7.2 + 50mM Mannitol added freshly) was prepared. 25μl of cell suspension mix with DNA was added into each well of the Nucleofection strip. Pulse programme DN-100 was used to deliver nucleofection pulse to each well. Control wells did receive a mock pulse without delivery of actual voltage.

### Cell culture of human and rodent lines

Human HEK293T cells were cultured in DMEM-F12 + 10% FBS + 1% Penicillin/Streptomycin. Murine NIH3T3 cells were cultured in DMEM + 10% FBS + 1% Penicillin/Streptomycin. The hamster ovary-derived cell line CHO-K1 was cultured in DMEM-F12 + 10% FBS + 1% Penicillin/Streptomycin. All lines were cultured at 37°C and 5% CO2 in a humidified incubator and regularly checked for mycoplasma contamination.

### sLCR activity transfection screening

Three cell lines of variable species-background were plated at a density of 3,000 cells (293T and CHO-K1) or 5,000 cells (NIH3T3) in their respective medium in black-walled 96-well-plates for optical imaging (Greiner, #655090). For transfection on the following day, we used the Fugene HD reagent and determined optimised conditions according to the manufacturers protocol for each cell line in a 96-well-plate format in a pre-experiment. In brief, we found 100ng of DNA and varying ratios of Fugene:DNA ratios (293T 2.5:1; NIH3T3 4:1; CHO-K1 4:1) to yield sufficient transfection efficiencies in a total reaction volume of 100μl per well. Mastermixes from 28 sLCR and three transfection control plasmids with Fugene reagent and serum-free RPMI were prepared accordingly and transferred to the screening plates in triplicates. After 48h of incubation time, nuclei were stained for 4h with 2μM Hoechst 33258 and fluorescent live-cell imaging for Hoechst, GFP, mCherry and iRFP was conducted on a high-content imaging platform (Operetta CLS, Perkin Elmer). We used the non-confocal mode and a 10x air objective to image the whole field of view of each well under live-cell imaging mode with temperature and CO2 control. LED power and detector exposure time were adjusted on non-sLCR transfection controls and untransfected wells.

### High-content screening analysis for sLCR activities

After filtering each fluorescent channel (sliding parabola 10px), we used the Harmony-Software building blocks to identify and count nuclei based on Hoechst-staining. Fluorescent cell-objects were identified based on GFP, mCherry or iRFP intensities. Fluorescent objects were filtered by applying a threshold for object size and mean intensities as well as number of objects were determined. Data with all relevant parameters was exported as csv files and analysed using R Studio. To account for differences in transfection efficiencies and to allow cross-comparison of sLCR expression among the three cell lines, we first calculated a transfection score as a proxy for efficiency. From the three non-sLCR transfection control plasmids (pMAX-GFP, UBC-mCherry, piRFP670) we established this score separately per cell line by determining the transfection_score = (control_fluorescent_objects_number / nuclei_number) for each control plasmid and calculated the combined mean from pMAX-GFP, UBC-mCherry and piRFP670. The value of this score represents the highest fluorescence intensity in the screen for each line and allows scaling of the sLCR plasmid activities between a value of 0 for untransfected controls and 1, as outlined in the following sentence. To assess the sLCR activity in each line, we calculated the sLCR_activity_score = (sLCR_fluorescent_objects_number / nuclei_number) and normalised this value by dividing through the previously established transfection score, that is setting the upper bar for the highest rate of fluorescent activity in each line and allows comparing sLCR activities across cell lines. Mean values of activity scores for each of the 28 sLCRs was calculated from the biological triplicates and data was plotted as boxplots using the ggplot2 package. Statistical testing was done through two-way ANOVA with multiple comparisons testing and Dunnett contrasts p-value adjustment.

### FACS sorting

Transduced/Transfected hGICs were harvested into single cell suspensions and resuspended into cold RHB-A complete medium and filtered into FACS tubes. Sorting was conducted using BD FACS Aria III or Fusion. The appropriate laser-filter combinations were chosen depending on the fluorophores being sorted for. Typically, to remove dead cells, events were first gated on the basis of shape and granularity (FSC-A vs. SSC-A) and doublets were excluded (FSC-A vs. FSC-H). Positive gates were established on PGK-driven and constitutively expressed H2B-CFP as sorting reporter (in case of Lentiviral construct) or iRFP expression (for PiggyBac-transfected cells).

### FACS analysis of MGT#1 and MGT#4 upon ME-GBM specific activators

Transduced/Transfected and sorted hGICs harbouring either Lentivirus-MGT#1 or PiggyBac-MGT#1/4 were seeded as single cells into 6-well plates in RHB-A medium supplemented with 10ng/ml TNFa (R&D Systems, 210-TA-020) or without TNFa as control. After 48h culture, hGICs were harvested into single cell suspensions and resuspended into cold RHB-A complete medium and filtered into FACS tubes. Events were first gated on the basis of shape and granularity (FSC-A vs. SSC-A) and doublets were excluded (FSC-A vs. FSC-H), then appropriate laser-filter combinations to analyse sLCR expression of mVenus, EGFP or mCherry were chosen. Data was analysed and visualized using FlowJo_v10.

### Genome wide CRISPRa library amplification, infection and library construction

For the genome-wide pooled CRISPR activation screen, we utilized the Human Genome-wide CRISPRa-v2 Library (Addgene Pooled Libraries #83978) consisting of 104,540 sgRNAs targeting 18,915 genes (top 5 sgRNAs per gene). Amplification of the library was performed according the reported Addgene protocol. Viral production was conducted in 293T cells, with viral titration performed before target cell transduction. To achieve a library representation over 240×, we transduced a total of 5 × 10^7^ of human glioma initiating cells harbouring the dCas9-VPR system as well as mesenchymal or proneural GBM reporters (IDH-wt-hGICs MGT#1, IDH-wt-hGICs-MGT#4, IDH-wt-hGICs-PNGT#3) at a multiplicity of infection of ∼0.5. The cells were kept for 15 days corresponding to 7 passages, after which the cells were FACS sorted for high and low reporter signal.The genomic DNA was extracted by lysing the cell pellets for 10 minutes at 56°C in AL buffer (Qiagen, 19075), supplemented with Proteinase K (Invitrogen, AM2548) and RNAse A (Thermo Scientific, 10753721), subsequently purified with AMPure XP beads (Beckman Coulter, A63881), and eluted in EB buffer (Qiagen, 19086). Next-generation sequencing (NGS) libraries were constructed in a two-step PCR setup, where the PCR1 is used to amplify the sgRNA scaffold, while the PCR2 introduced Illumina compatible adaptors with unique P5/P7 barcodes, allowing sample multiplexity. For the PCR1, each gDNA sample was divided over 20 to 26 reactions, which were subsequently pooled together and purified using AMPure XP beads. The optimal cycle numbers for PCR2 were determined for 1 μL of each PCR1 individually by conducting a qPCR amplification using KAPA HiFi HotStart Ready Mix (Roche, 7958927001) and 1× EvaGreen (Biotium, 31000). 5 ng of the purified PCR1 of each sample was used as input for the final PCR2. Both PCR1 and PCR2 were performed using KAPA HiFi HotStart Ready Mix. Primers are available upon request. Quality control of the final libraries was performed using the Qubit dsDNA HS kit (Invitrogen, Q32854) and Collibri™ Library Quantification Kit (Invitrogen, A38524500) for quantification and TapeStation High Sensitivity D1000 ScreenTapes (Agilent, 5067-5584) for determination of PCR fragment size. The barcoded libraries were pooled together in equal molarities and sequenced on an Illumina NextSeq500 using the 150 cycles V2.5 chemistry (1×150 nt single-read mode). The 20 bp gRNA sequences were extracted using a custom script and aligned to the Human Genome-wide CRISPRa-v2 Library reference using bwa to subsequently generate the gRNA read counts. For assessing CRISPRa hits, relevant for the utilized model system, the gRNA counts were filtered for targets denoted by high expression in the IDH-wt-hGICs (CPM > 5). Differential gRNA abundance and candidate targets prediction was conducted using the gCrisprTools R package. Further, to identify GBM context-specific CRISPRa hits, we performed orthogonal convergence of TCGA differential expression (TCGA GBM, Verhaak et al., ^29^) together with the glioblastoma stem cells state-specific dependencies (Richards et al., ^30^). For follow up and subsequent validation we prioritized hits, that characterized by high TCGA GBM patients subtype specific expression (LFC > |2|), enrichment in glioblastoma stem-cells CRISPRout genome-wide screening (z-scores >|2|) and median log2FC > 0 and Rho enrichment score < 0.5 in our LSD-sLCR CRISPRa screens. Ingenuity pathway analysis used as input gCrisprTools-derived fold-changes and FDR values, and ranked canonical pathways and upstream regulators by Fisher’s exact test.

---

**Figure S1.**
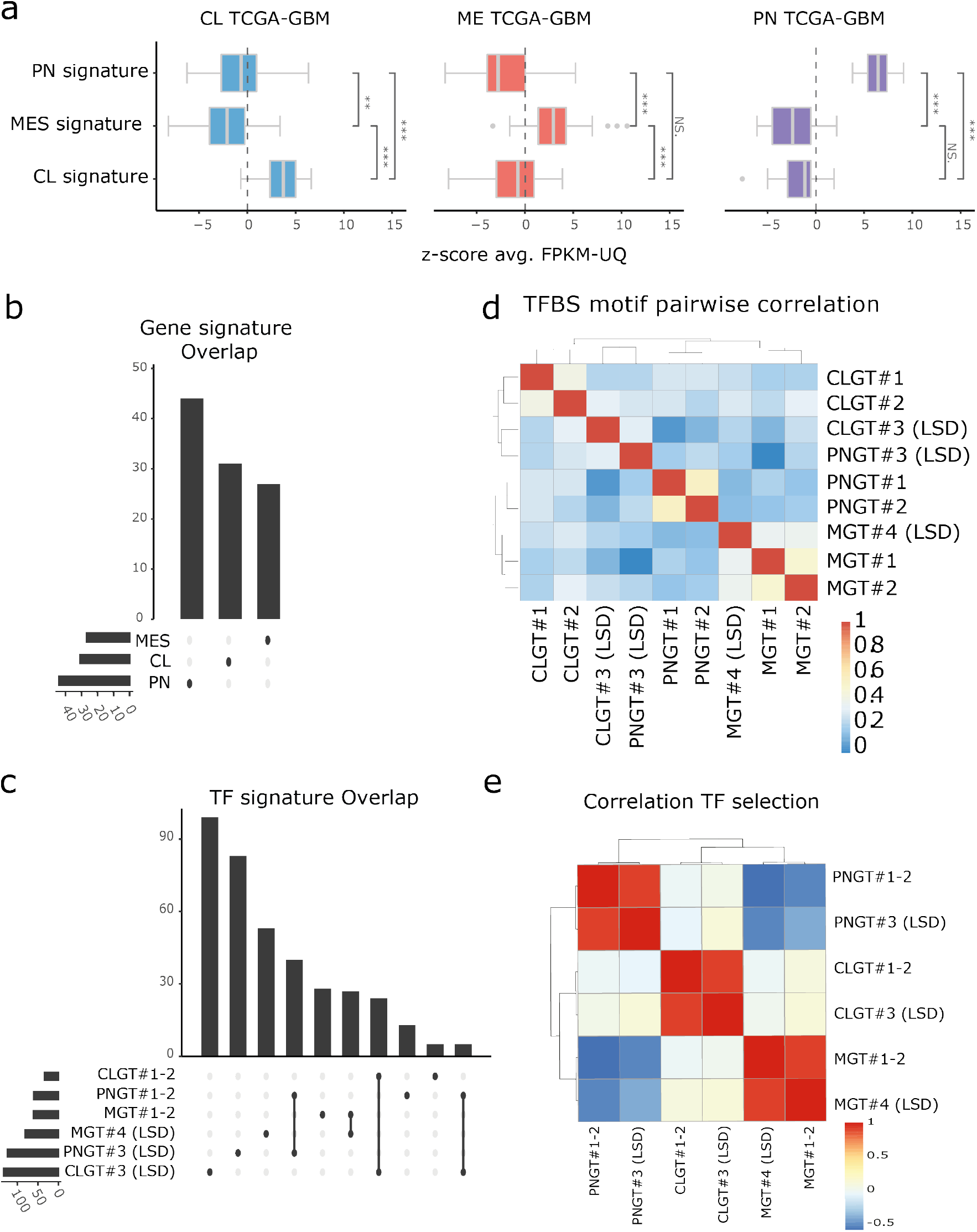
Correlation analyses of LSD input signature and TF lists. a) Boxplot comparing gene expression z-scores for the indicated sLCR’s signature gene sets in glioblastoma patients (TCGA-GBM, FPKM-UQ). Each annotated GBM transcriptional subtype features one comparison by Wilcoxon test (padj<0.05). b) Upset plot depicting the number of and overlap between signatures genes underlying each 1st generation/LSD GBM subtype sLCR. c) Upset plot depicting the number of and the overlap between TF underlying each GBM subtype sLCR designed by LSD. The connected lines denote the overlaps. d) Heatmap showing the Pearson correlation between the indicated input TFBS lists with respect to overall TFBS. Hierarchical clustering analyses used Euclidean distance and complete linkage. e) Heatmap showing the Pearson correlation of ssGSEA enrichment scores in Glioblastoma patients’ expression for each TF input lists. Both hierarchical clustering analyses used Euclidean distance and complete linkage.

**Figure S2.**
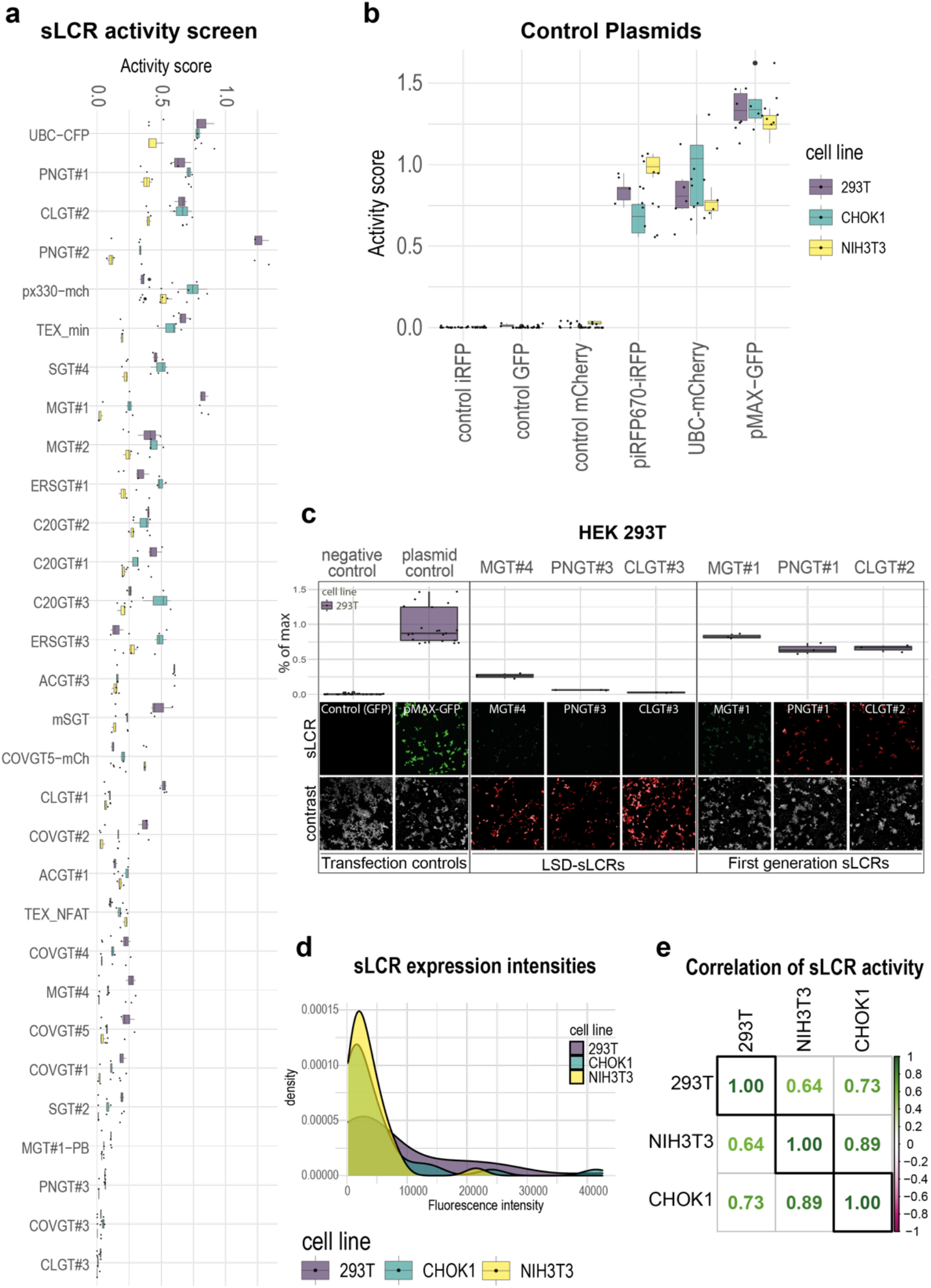
LSD allows designing functional and specific sLCRs. a) Box plot of indicated sLCRs (n=28) transfected in human epithelioid 293T, hamster epithelioid CHO-K1 and mouse fibroblastoid NIH3T3 cell lines. The top axis shows fluorescence normalized by controls and transfection efficiency per cell line. Each sLCR measurement was assessed in technical replica (n=3). b) Activity score normalisation with untransfected negative controls (n=9) and positive transfection control plasmids CFP, GFP, mCherry and iRFP670 (n=6), with expression driven by non-sLCR promoters. c) Upper, box plot comparison of activity scores for LSD and first generation sLCRs. Below, fluorescence microscopy images of sLCR expression (top panel) and contrast channel of either brightfield or independent promoter-driven fluorophore (lower panel). d) Density plot of raw fluorescence intensities of all datapoints for all sLCRs (n=28) in 293T, CHO-K1 and NIH3T3 cells. e) Pairwise pearson-correlation matrix of human and non-human cell lines for calculated sLCR (n=28) activities.

**Figure S3.**
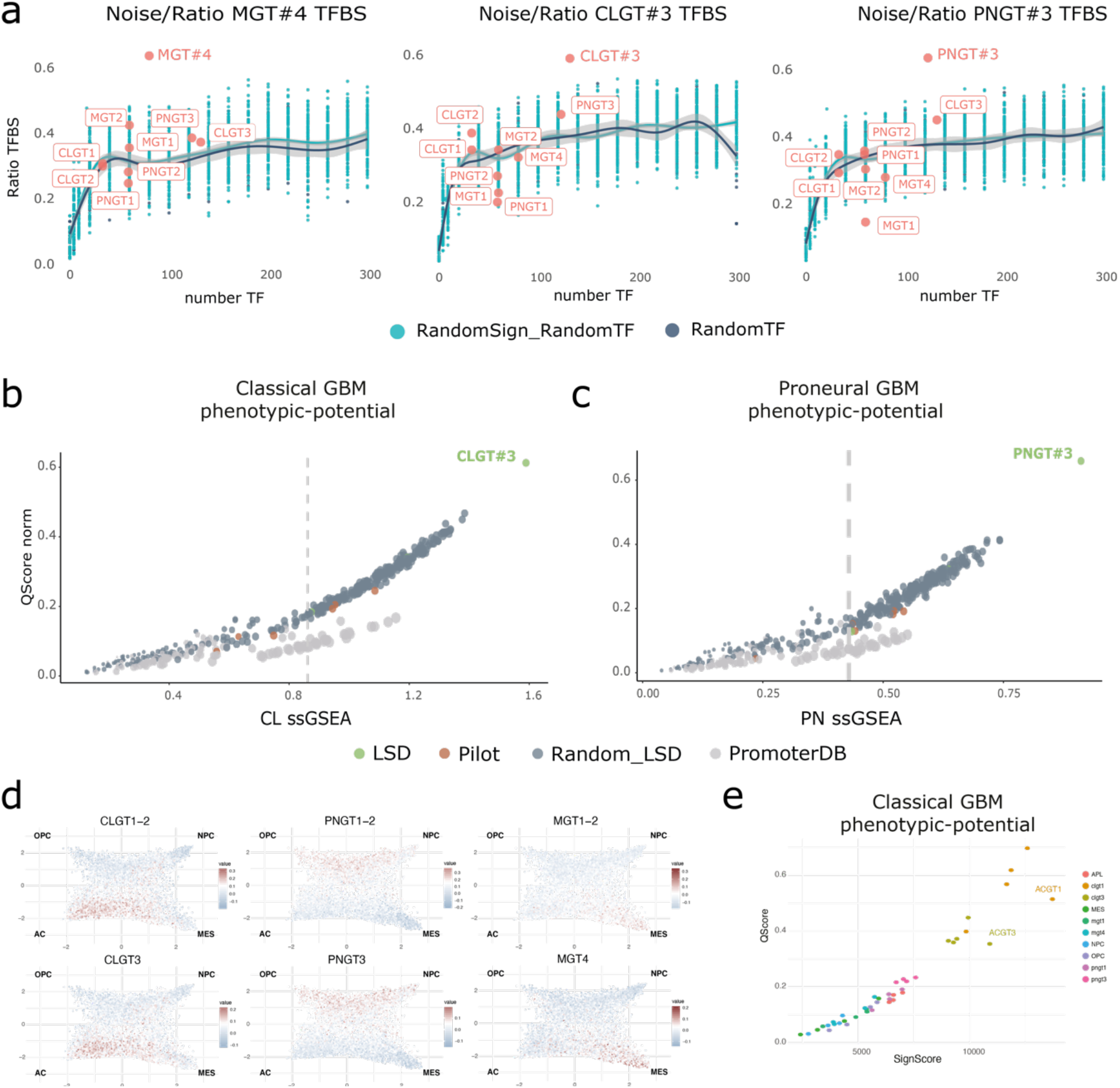
Extended analysis of LSD sLCRs phenotypic potential. a) Scatter plot showing the TFBS affinity ratio for on-target, off-target and randomly selected sLCRs. The Y-axis indicates the observed/expected ratio (i.e. observed/input TFBS) for the TFBS lists indicated in the header. The X-axis denotes the number of non-redundant input TF (see Methods). First-generation and LSD-designed sLCR are indicated. sLCRs were designed using LSD and input from random sampling of TFBS on mesenchymal GBM signature genes (random TF) or random signature genes (random Sign-TF). Fitted lines indicate LOESS regression. b-c) Scatter plot showing the signature score (x-axis) and affinity score (y-axis; see Methods) of the indicated reporters for the classical and proneural phenotypes. d) Comparison of ssGSEA normalized scores for signature genes for the indicated first generation sLCRs (above) and LSD-derived sLCRs (below). The cell states identified by Neftel et al. 2019 are indicated in each quadrant, and the original single-cell position is maintained in the two-dimensional representation. e) Scatter plot showing the signature score (x-axis) and affinity score (y-axis; see Methods) of the indicated reporters for the classical phenotype.

**Figure S4.**
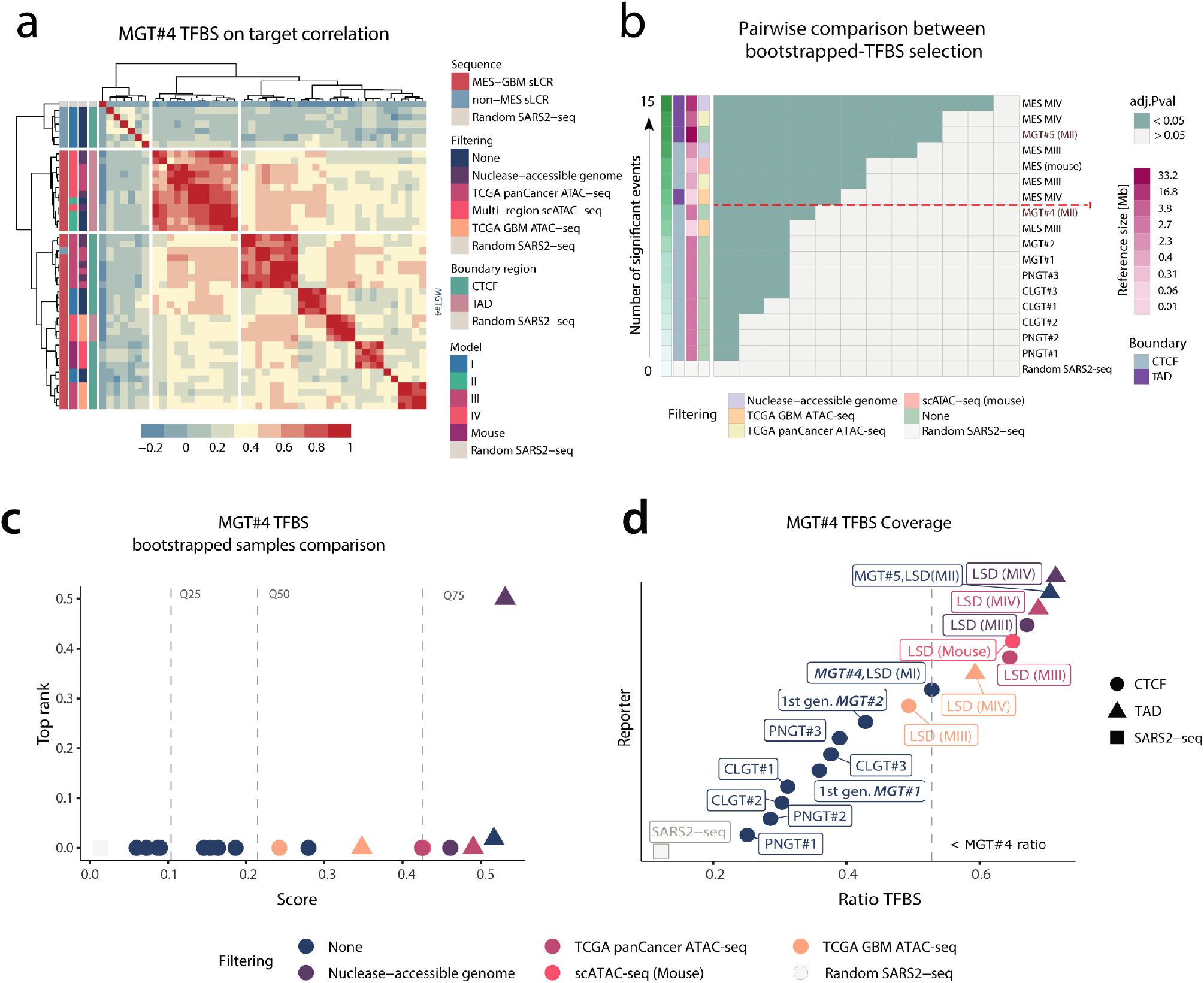
Extended analysis of mesenchymal GBM sLCRs generation with custom inputs. a) Heatmap showing Pearson correlation between the MGT#4 TFBS score of the indicated sLCRs. Hierarchical clustering used Manhattan distance. b) Heatmap showing the significance of pairwise comparisons between the indicated sLCRs after bootstrapping of the MGT#4 TFBS input (see methods). Color-code annotations denote the number of significant pairwise correlations between distributions (adj.Pval <0.05), the boundary annotation method (i.e. nearest-neighbour CTCF or annotated TADs), the type and size of reference genome/subset used for TFBS mapping. The color-coding for the size of the genome input is also indicated. Dotted line marks threshold for models with improved number of significant events. c) Scatter plot visualisation of the MGT#4 TFBS bootstrapped samples ranking based on the number of significant pairwise correlations between distributions (see Methods). d) Scatter plot visualisation of MGT#4 TFBS coverage. Axis represent the observed/expected TFBS ratio (x-axis) of the indicated reporters (y-axis). Dashed lines highlights the threshold of MGT#4 TFBS ratio.

**Figure S5.**
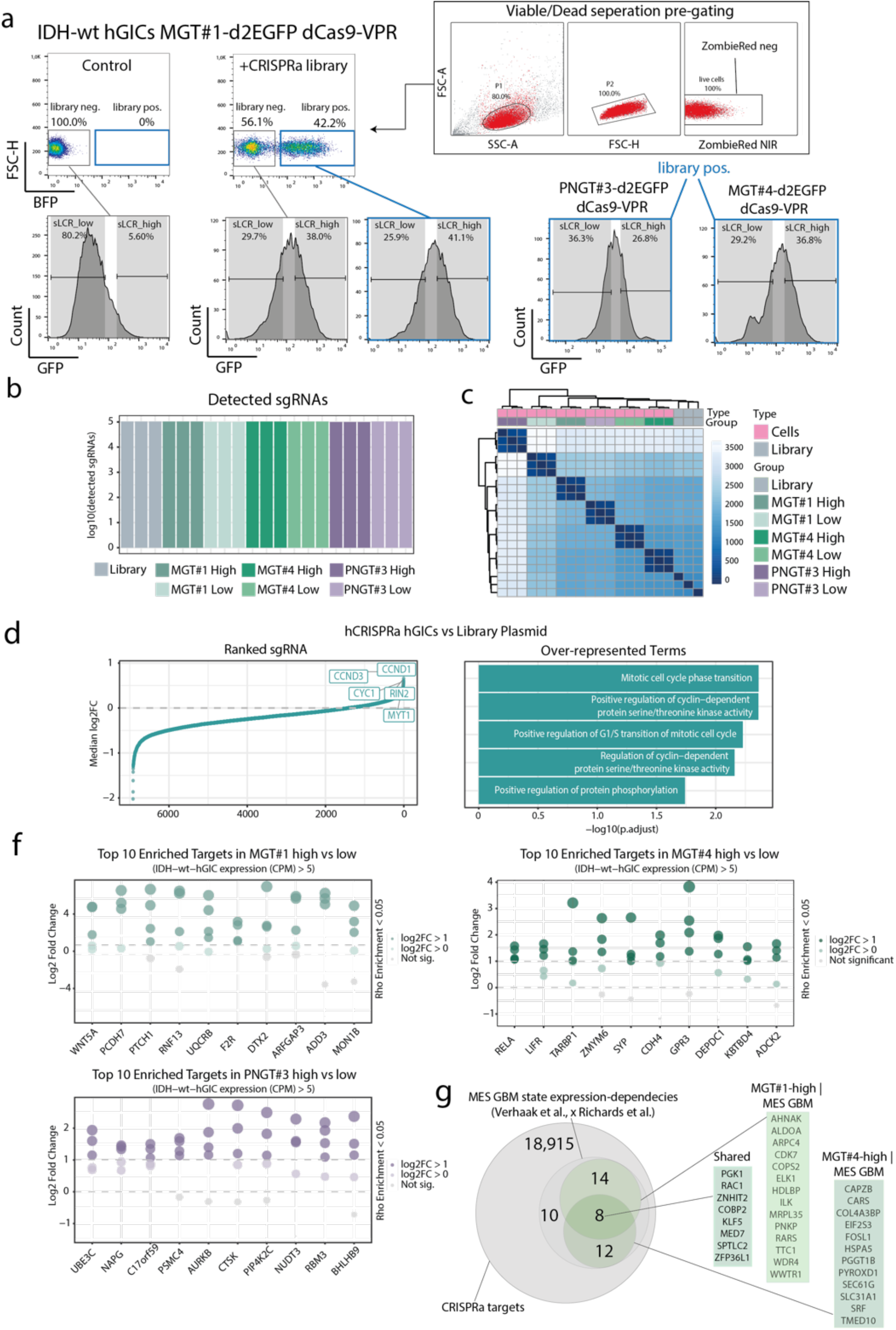
Extended analysis of the genome-wide CRISPR activation screens. a) FACS gating strategy for sLCRs and hCRISPRa-v2. Note that BFP-cells (non-transduced by the CRISPRa gRNA library) do also show similar levels of reporter expression, indicating that non-autonomous mesenchymal transition occurred. b) Barplot showing gRNA read counts across the indicated conditions. Homogeneous counts support library representation to be qualitatively maintained. c) Heatmap showing the Euclidean distance of normalized gRNA abundance in the indicated conditions. Color legend highlights sample origin (type) or level of reporter expression (group). d) Median log2-fold change based ranking of differentially abundant gRNAs between cells and library samples. e) Gene Ontology terms associated with the upregulated gRNA targets (median log2-fold change > 0.5) from data in (d). f) Dot plot visualisation of the top gRNA targets enriched in the indicated high reporter expressing cells. Dot color and size denote significance for each comparison and dashed lines denotes the indicated thresholds. g) Venn diagram showing overlap between the indicated datasets. Private and common candidate gene drivers from MGT#1-high and MGT#4-high screens are reported to the right.

